# Emergence of enteroaggregative *Escherichia coli* within the ST131 lineage as a cause of extraintestinal infections

**DOI:** 10.1101/435941

**Authors:** Erik J. Boll, Marc Stegger, Henrik Hasman, Louise Roer, Søren Overballe-Petersen, Kim Ng, Flemming Scheutz, Anette M. Hammerum, Arnold Dungu, Frank Hansen, Berit Lilje, Dennis Schrøder Hansen, Karen A. Krogfelt, Lance B. Price, James R. Johnson, Carsten Struve, Bente Olesen

## Abstract

*Escherichia coli* sequence type 131 (ST131) is a major cause of urinary and bloodstream infections and its association with extended-spectrum β-lactamases (ESBL) significantly complicates treatment. Most notorious is its rapidly expanding *H*30-Rx clade (named for containing allele 30 of the type-1 fimbrial adhesin gene *fimH* and extensive antimicrobial resistance), which appears to have emerged in the United States due in part due to the acquisition of the ESBL-encoding *bla*_CTX-M-15_ gene and resistance to fluoroquinolones. However, non-*H*30 ST131 lineages with acquired CTX-M-type resistance genes also are emerging. Based on whole-genome analyses, we describe here the presence of an (*fimH*) *H*27 *E. coli* ST131 lineage that currently is causing an outbreak of community-acquired bacteremia and recurrent urinary tract infections (UTIs) in Denmark. This lineage has acquired both a virulence plasmid (pAA) that defines the enteroaggregative *E. coli* (EAEC) diarrheagenic pathotype and multiple genes associated with extraintestinal *E. coli* (ExPEC) that combined has made this particular ST131 lineage highly successful at colonizing its human host and cause recurrent UTI. Moreover, using a historic World Health Organization *E. coli* collection and publically available genome sequences, we identify a global *H*27 EAEC ST131 lineage dating back as far as 1998. Most *H*27 EAEC ST131 isolates harbor pAA or pAA-like plasmids, which analysis strongly imply was caused by a single ancestral acquisition. These findings illustrate the profound plasticity of this important pathogenic *E. coli H*27 lineage in general, and the genetic acquisitions of EAEC-specific virulence traits that likely confer an enhanced ability to cause intestinal colonization.

**Importance:** The *E. coli* ST131 lineage is a notorious extraintestinal pathogen. A signature characteristic of ST131 is its ability to asymptomatically colonize the gastrointestinal tract and then opportunistically cause extraintestinal infections, such as cystitis, pyelonephritis and urosepsis. In this study, we report a novel ST131 sublineage that has acquired the enteroaggregative diarrheagenic phenotype, spread across multiple continents and has been associated with multiple outbreaks of community-acquired bloodstream infections in Denmark. The strain’s ability to both cause diarrhea and colonize the human gastrointestinal tract may facilitate its dissemination and establishment in the community, whereas the strain’s clonal nature may facilitate targeted control strategies, such as vaccination.

## Introduction

*Escherichia coli* sequence type 131 (ST131) is the dominant multidrug-resistant (MDR) extraintestinal pathogenic *E. coli* (ExPEC) lineage worldwide, causing a wide range of infections including bloodstream and urinary tract infections (BSIs/UTIs) (*1–3*). Its rise to global dominance and pathogenicity is thought to have been primed by sequential acquisition of virulence-associated genes followed by development of antibiotic resistance (*4, 5*). ST131 predominantly exhibits serotype O25:H4 and is closely associated with fluoroquinolone resistance and the production of the CTX-M-15 extended-spectrum β-lactamase (ESBL) (*2, 6*). The expansion of *E. coli* ST131 in the United States has been driven mainly by a single clade, designated *H*30 because of its tight association with allele 30 of the type-1 fimbrial adhesin gene, *fimH*. *H*30 has a prominent MDR-associated clade, *H*30R, which accounts for most fluoroquinolone resistance within ST131. *H*30R in turn has two main sub-clades: *H*30R, which accounts for almost all ST131-associated CTX-M-14 and CTX-M-27 production (although most members are ESBL-negative), and *H*30Rx, which accounts for almost all ST131-associated CTX-M-15 production (*6*). In addition to *H*30, other distinct clades of ST131 are also circulating worldwide, most commonly associated to the 22 and 41 *fimH* alleles. While these non-*H*30 allelic variants are normally not associated with carriage of CTX-M-genes, cases of *bla*_CTX-M-14_, *bla*_CTX-M-15_ and *bla*_CTX-M-27_ acquisition by ST131 lineages carrying *fimH*22 or -*H*41 have been reported (*4, 7*).

ST131 isolates typically carry multiple ExPEC-associated virulence genes encoding adhesins, toxins, and siderophores, whereas virulence genes typical of diarrheagenic *E. coli* (DEC) rarely have been reported (*8, 9*). However, we recently surveyed an international World Health Organization (WHO) collection of historic *E. coli* for ST131 O25 isolates, assessing them for temporal trends of antibiotic resistance and virulence traits. Among a total of 128 ST131 isolates we found 12 (9%) that fulfilled molecular criteria for the enteroaggregative *E. coli* (EAEC) pathotype (*10*). Pulsed-field gel electrophoresis (PFGE) analysis revealed a cluster comprising seven of these EAEC isolates. Of these, six - including two urine isolates from patients with UTI, three fecal isolates from patients with diarrhea, and one lower respiratory tract isolate (associated symptoms unknown) - were from Danish patients (1998-2000), supporting the occurrence of an unrecognized EAEC ST131-associated outbreak of UTI and possibly also diarrhea in this time period in Denmark (*10*).

EAEC strains cause endemic diarrheal illness in developing countries and foodborne outbreaks in developed countries, and have been associated with extraintestinal infections (*11*). This pathotype gained particular attention following a major foodborne outbreak in Germany in 2011 that was caused by a Shiga toxin (Stx)-producing O104:H4 EAEC strain and resulted in 3,842 confirmed cases and 54 deaths (*12*). Additionally, a worryingly high prevalence of multidrug resistance among EAEC strains has been reported, and reports of CTX-M-type ESBL-producing EAEC have appeared recently from around the world (*13–16*). EAEC pathogenesis involves adherence to human intestinal mucosa by virtue of aggregative adherence fimbriae (AAF) and subsequent biofilm formation. The AAF are encoded on large plasmids designated pAA that also encodes a suite of other EAEC virulence factors, including AggR, a global regulator of EAEC virulence; dispersin, required for proper dispersal of AAFs on the bacterial surface; the AatPABCD transporter system, which mediates dispersin secretion; and Aar, a recently described negative regulator of AggR (*17, 18*).

To understand the emergence and underlying genetic acquisitions leading to ESBL-producing EAEC ST131, we investigated the relatedness of a large collection of temporally and spatially diverse ST131 isolates. This investigation revealed the emergence of a global *H*27 lineage of EAEC linked to a single acquisition of the pAA plasmid that likely improved the strain’s ability to persistently colonize its human host.

## Results

### Identification of historic Danish EAEC ST131

Our previous PFGE-based analysis of ST131 O25 isolates within the WHO *E. coli* collection identified an apparently unrecognized outbreak of UTI, and possibly also diarrhea, in Denmark in 1998-2000 that was caused by a group of seemingly similar EAEC isolates (*10*). To better estimate these isolates’ relatedness and to correct for recombination (*6*), all 128 ST131 isolates from the WHO collection were subjected to whole-genome sequencing. The *in silico* analyses regarding the genotypic features of the isolates revealed that the most frequent *fimH* alleles were *H*30 (51%, n=65), *H*22 (34%, n=44) and *H*27 (7%, n=9). Most of the isolates harbored multiple antibiotic resistance genes. The most common ESBL gene was *bla*_CTX-M-15_ (37%, n=47), which occurred almost exclusively within the *H*30 subgroup (98%, 46/47). See **Suppl. Table 1.**

To ensure that only high-quality genome data sets were included in our SNP analysis, we excluded the genome data for seven (5%) of the 128 ST131 isolates due to low sequencing depth. In addition, we included genome sequences from a published collection of human clinical ST131 *E. coli* (n = 93) collected in the USA and Germany between 2010 and 2012 (*6*). Across these genomes, 11,529 SNPs were identified within ~38.4% of the JJ1886 genome that was conserved across all isolates. After purging of recombinant regions, 4,241 SNPs were used to infer relationships between the isolates.

The SNP-based phylogeny showed distinct overall clustering of isolates in accordance to their *fimH* alleles, with all 116 isolates carrying the *fimH*30 allele clustering as a monophyletic clade (**Suppl. Fig. 1**). Likewise, all 10 isolates carrying the *fimH*27 allele clustered within a single clade which - like the *H*30 clade - appeared to descend from an ancestral *H*22 group. Of these *fimH*27-carrying isolates, eight - all from the WHO collection - qualified molecularly as EAEC based on presence of ≥1 of the EAEC-associated putative virulence genes *aggR*, *aatA*, and *aaiC (19)*. On the basis of previous PFGE data (*10*), seven of these eight EAEC isolates had formed a cluster, whereas the remaining isolate (C796-00) appeared quite distinct from the others. By contrast, here all eight isolates clustered together, with C796-00 being most closely related to three of the other seven EAEC isolates (**Suppl. Fig. 1**). Included in this cluster of *fimH*27-carrying EAEC isolates were C86-04, a fecal isolate from Vietnam (2004), and seven Danish isolates, including two urine isolates, one lower respiratory tract isolate, and four fecal isolates from patients with diarrhea, suggesting that this lineage was able to cause both extraintestinal infections and diarrhea.

**Fig. 1.**
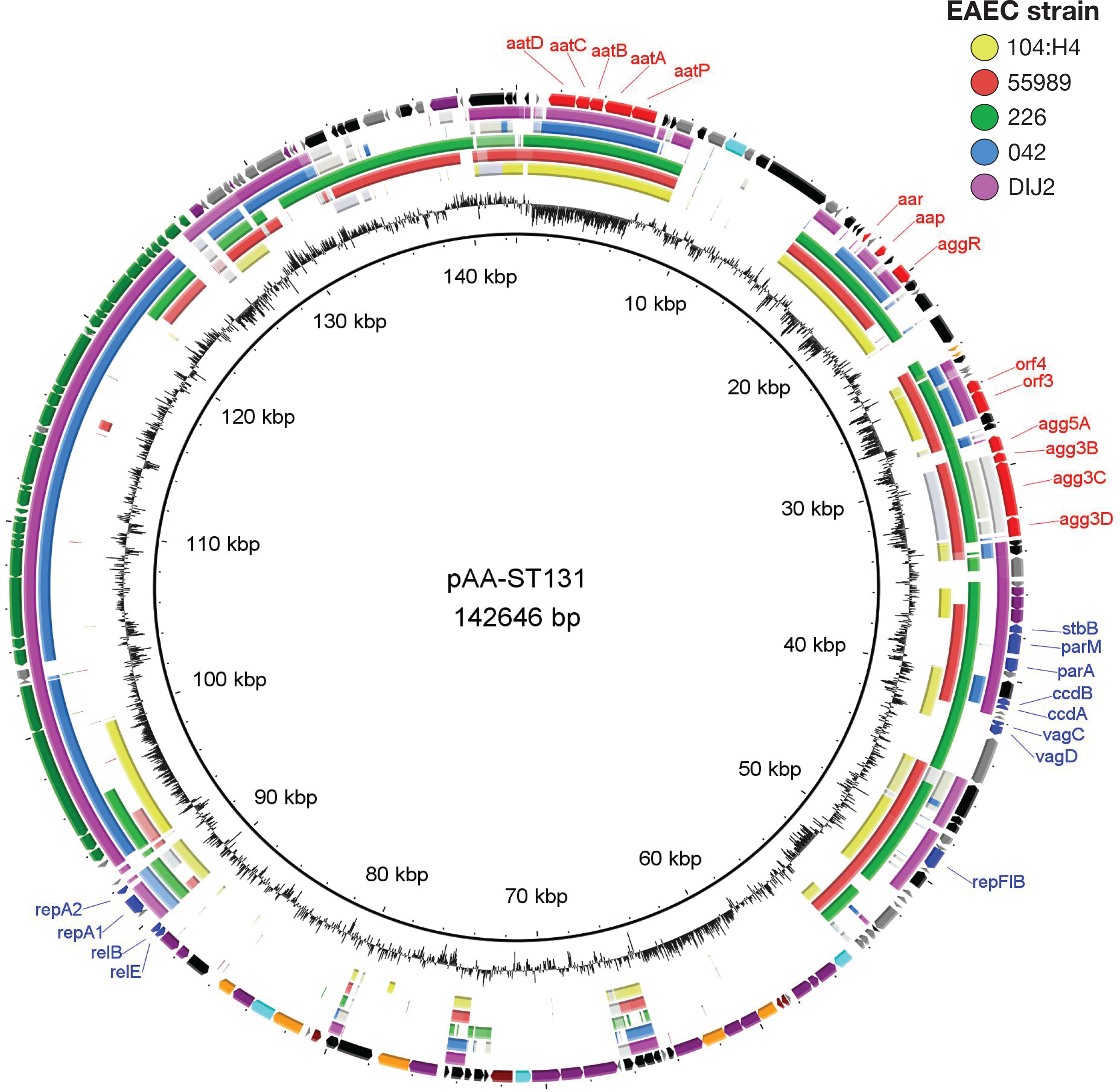
Circular map of plasmid pAA-ST131 compared to other publicly available pAA plasmids. The outer ring shows predicted ORFs of pAA-ST131. Colors represent different putative functions: gray, hypothetical proteins; red, EAEC-specific virulence factors; blue, plasmid replication and maintenance; maroon, catabolism and metabolism; orange, membrane and transporter proteins; green, conjugational transfer proteins (*tra* and *trb* genes); light blue, regulatory genes; purple, miscellaneous; and black, mobile elements. Within the circles representing pAA plasmids from other EAEC strains (labeled one to five), the darkest color indicates >90% nucleotide identity, the lightest color >80% identity.

Unlike the PFGE analysis, the SNP-based analysis additionally revealed a distinct cluster of isolates located within the *H*22 group consisting of the remaining four EAEC isolates from the WHO collection (**Suppl. Fig. 1**). Three of the isolates carried *fimH*22, whereas one isolate carried *fimH*298, which differs from *fimH*22 by only a single nucleotide. All four isolates were from urine samples collected within a 10-month period in 1998, from elderly patients admitted to four different departments of the same hospital in the Capital Region in Denmark, suggesting an unrecognized nosocomial UTI outbreak. With the exception of Vietnamese isolate C86-04, all 11 EAEC isolates were recovered from Danish patients. The Vietnamese isolate carried *bla*_CTX-M-27_, whereas no ESBL genes were present in the other 11 EAEC isolates. (**Suppl. Table 1**).

### EAEC virulence genes in historic Danish EAEC ST131

The four clustered EAEC urine isolates that carried *fimH*22 or *fimH*298 all harbored the *aggDCBA* gene cluster encoding the AAF/I variant, whereas seven of the eight clustered *fimH27*-carrying EAEC isolates of diverse sources all harbored *agg5DCBA* encoding the recently described AAF/V variant (*20*). The remaining *H*27 isolate, C1883-99, lacked the *agg5A* gene (**Suppl. Table 2**). All but two of the 12 EAEC isolates contained genes encoding AggR, Aar, dispersin (*aap*), and the AatPABCD transporter system, plus two additional AggR-regulated open reading frames (ORFs), ORF3 and ORF4, that are assumed to play a role in isoprenoid biosynthesis (*21*). The remaining two isolates, C1883-99 (*H*27) and C167-00 (*H*22), both lacked most of these EAEC-specific genes, implying partial deletion of the pAA plasmid in isolates from both these *fimH*-associated clades.

### Invasive EAEC ST131 isolates with *bla*_CTX-M-101_ obtained from Danish patients

To determine whether EAEC ST131 strains were present among contemporary Danish patients, we searched for EAEC-specific virulence genes within a recently published collection of 552 whole-genome sequenced ESBL-producing *E. coli* (ESBL-*Ec*) isolates obtained from patients with BSIs in Denmark between 2014 and 2015 (*22*). ST131 accounted for 50% (n = 258) of the isolates (*22*). Among these, 25 isolates harbored the genes encoding AAF/V, AggR, Aar, Aap, AatPABCD, and ORF3/4 (**Suppl. Table 2**). All 25 carried *fimH*27 and *bla*_CTX-M-101_. In total, 27 isolates in the collection were *bla*_CTX-M-101_-positive (*10*), of which thus 93% qualified molecularly as EAEC. The EAEC pathotype did not occur in conjunction with any other ESBL-encoding gene; hence, the association between *bla*_CTX-M-101_ and the EAEC pathotype was highly significant (χ^2^, p<0.0001), suggesting that the resistance gene may be co-localized with EAEC-specific virulence genes on the pAA plasmid. As shown by Roer *et al.*, the 27 ST131 strains with *bla*_CTX-M-101_ formed a distinct cluster with nine or fewer SNP differences, supporting a recent emergence (*22*).

To assess whether the same ESBL-EAEC ST131 strain could be detected in the urine of the patients from whom the positive blood samples were collected, urine isolates from eight of the 258 patients with an ESBL-EAEC ST131 BSI were available and sequenced. Notably, all of these patients had presented with UTI either prior to, concurrent with, or subsequent to the time of blood sampling. Thus, some patients provided more than one urine sample. Specifically, one patient presented 19 days after an initial bacteremia episode with both recurrent bacteremia and UTI, all involving the ESBL-EAEC ST131 strain, and four other patients had recurrent same-strain UTI diagnosed from three weeks to eight months after their initial UTI episode. SNP analysis demonstrated that the corresponding blood and urine isolates were nearly identical (i.e., differed by ≤6 SNPs), whereas different pairs were separated by zero to 11 SNPs. Of the eight pairs, five clustered distinctively in the phylogeny thus strongly implying their relatedness. For the remaining three pairs of blood and urine isolates, the data were inconclusive (**Suppl. Fig. 2**). These findings suggest strongly that the ESBL-EAEC ST131 strains were capable of long-term persistence in these hosts and/or their immediate environment and of causing repeated episodes of UTI and bacteremia.

**Fig. 2.**
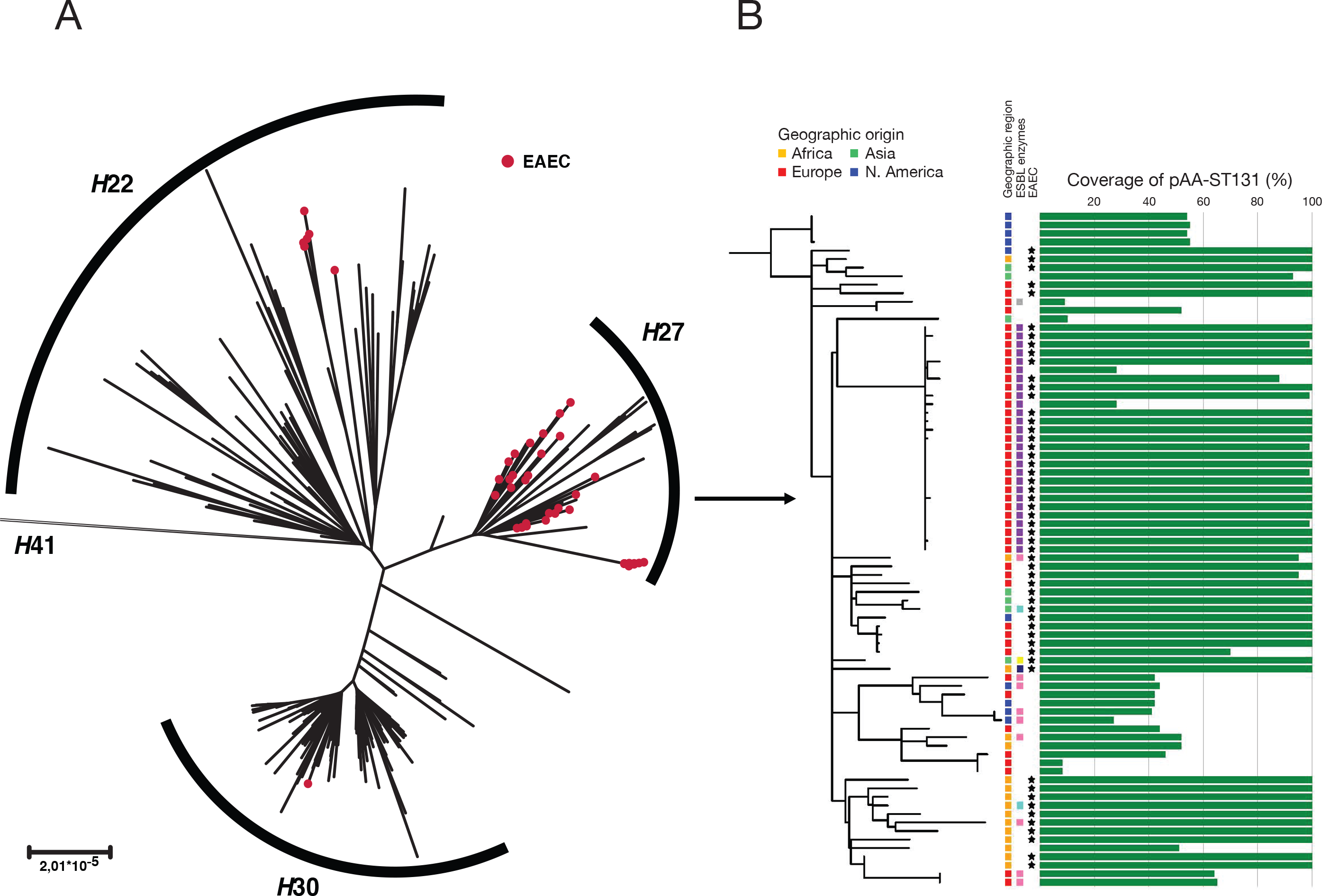
A) Unrooted phylogenetic tree of ST131 genomes from the WHO collection, the US/German collection from 2010-2012, the Danish *bla*_CTX-M-101_-containing isolates from 2014-2015, and the EnteroBase database of international isolates carrying *fimH*27 (n=287). The distant *H*41 cluster is not shown. EAEC isolates highlighted in red. B) Rooted phylogenetic tree with all isolates within the *H*27 clade (n=79). ESBL-enzymes: CARB-2 (grey), CTX-M-101 (purple), CTX-M15 (pink), CTX-M-15+OXA-10 (dark blue), CTX-M-27 (yellow) and SHV-12 (light blue). EAEC-positive isolates are marked by an asterisk. Scale bar represents substitution rate in the conserved core genome.

### pAA plasmid characterization in ST131 isolates

To characterize in detail the pAA plasmids present in the ESBL-EAEC ST131 strains, we applied MinION sequencing (Oxford Nanopore Technologies) to obtain the complete plasmid sequence for one such plasmid, which we designated pAA-ST131, from a single representative Danish *E. coli* isolate (ESBL20150001, Sequence Read Archive (SRA) ID: DF215RRW). The complete plasmid sequence was 142,646 bp in length, with an average G+C content of 48.7% (**Fig. 1**, GenBank accession no. KY706108). A total of 197 open reading frames (ORFs) were predicted and annotated, of which 142 were functionally assigned. Two replicons were identified, RepFII and RepFIB, with the multireplicon F plasmid FAB formula of F1:A-:B33.

pAA-ST131 contained several genes associated with plasmid stability, including *parA*, *parM*, and *stbB*, plus three toxin-antitoxin (TA)-based addiction systems: *ccdAB*, *vagCD*, and *relBE* (**Fig. 1**). Furthermore, it harbored a 33 kb complete *tra* region encoding transfer components (24 *tra* genes, 8 *trb* genes, and *finO*), implying that the plasmid may be conjugally transferable. It also contained genes encoding putative transmembrane proteins and proteins involved in catabolism and metabolism, plus several integrated mobile elements. It contained all the EAEC virulence factor genes already identified by Illumina sequencing (as expected), but no additional known virulence factor genes or antibiotic resistance genes including *bla*_CTX-M-101_. Because isolate ESBL20150001 contains no additional plasmids (**Suppl. Fig. 3**), we conclude that *bla*_CTX-M-101_ is chromosomally located.

A BLAST comparison of pAA-ST131 with five complete pAA plasmids (Genbank accession IDs NC_018666 (O104:H4), NC_011752 (55989), FN554767 (042) and NC_008460 (DIJ1) and SRA ID SRA055981 (226)) showed that all these plasmids shared common features (**Fig. 1**), mostly corresponding to EAEC virulence genes. The entire genomic sequences were not available for all corresponding EAEC isolates but they belonged to at least three STs (ST40, ST414, and ST678). However, they also varied substantially for genetic content, as described previously for *E. coli* virulence plasmids (*23*). Specifically, regions containing putative metabolic, catabolic, and transmembrane proteins were unique to pAA-ST131 (**Fig. 1**). Additionally, although (like pAA-ST131) four of the five reference pAA plasmids also contained the RepFIB and/or the RepFIIA replicon, only three contained both. Moreover, only two contained the *tra* region.

### Global emergence of EAEC ST131 isolates carrying *fimH*27

To assess the global extent of EAEC ST131 isolates, we screened for *aggR* (the global regulator of EAEC virulence) among all >3,500 *E. coli* ST131 genomes (as of November 15^th^ 2017) available in EnteroBase (http://enterobase.warwick.ac.uk), a database with >100,000 genomic assemblies of enteric bacteria, including *E. coli*. Disregarding the *aggR*-positive, *bla*_CTX-M-101_-containing Danish EAEC isolates, we identified another 25 international *aggR*-positive isolates, which carried the following *fimH* alleles (no. of isolates, % of 25): *fimH*27 (18, 66%), *fimH*22 (one, 4%), *fimH*298 (two, 8%), *fimH*30 (one, 4%), *fim*H5 (two, 8%), and *fim*H54-like (one, 4%) (**Suppl. Table 2**). The *aggR* gene occurred in 18 (38%) of the 47 other *fimH*27-carrying ST131 isolates in EnteroBase.

To determine whether the 12 EAEC ST131 isolates from the WHO collection were clonally related to other available EAEC ST131 isolates, a SNP-based phylogeny was constructed using 1) all 121 ST131 isolates from the WHO collection (*10*); 2) 93 German/US ST131 isolates from 2010-2012 (*6*); 3) the 27 Danish *bla*_CTX-M-101_-containing ST131 isolates from 2014-2015 (*22*); and 4) 46 international *fimH*27-carrying ST131 isolates from Enterobase. Focusing first on isolates within the *H*22 clade (**Fig 2A**), WHO collection EAEC isolate C180-00, carrying *fimH298*, clustered together with two *fimH*298 EAEC EnteroBase isolates (SRA IDs SRR2970775 and SRR2970774) from Cambodia (2009-2010). In contrast, the three WHO collection EAEC isolates carrying *fimH*22 were not closely related to the only *fimH22* ST131 isolate from EnteroBase.

In the rooted phylogeny, most EAEC isolates nested within the *H*27 clade (**Fig. 2A**). To improve resolution, this clade’s 79 isolates (76 with *fimH27*, two with *fimH5*, one with *fimH54*) were analyzed separately. The 27 *bla*_CTX-M-101_-containing Danish EAEC isolates formed a distinct cluster with very short branches indicating a recent emergence (**Fig. 2B**). By contrast, none of the *fimH*27-carrying international (i.e., non-Danish) isolates clustered either with the Danish isolates or with one another, suggesting that the Danish outbreak was confined to Denmark, and that no other focal outbreaks were captured. The WHO collection’s *H*27 clade EAEC isolates were intermingled with isolates from the UK, Thailand, and Canada (**Fig. 2B**), suggesting global spread of a common-ancestry strain.

To determine the mosaicism of the EAEC-specific virulence genes among *fimH*27- carrying ST131 isolates, sequence reads from all 79 *H*27 clade isolates were mapped against pAA-ST131 from ESBL20150001. Intriguingly, pAA-ST131 was highly conserved among the *H*27 EAEC isolates (**Fig. 2B)**. To validate the *in silico* results, plasmid gel profiling was done for representative *fimH*27-carrying *bla*_CTX-M-101_-containing EAEC isolates and EAEC isolates from the WHO collection. This confirmed that all but three of the ten tested isolates contained a single plasmid of conserved size (~140 kb). When mapping the sequence reads to pAA-ST131 these all showed 99-100% coverage. The three exceptions included WHO isolate C1883-99, which harbored a slightly smaller plasmid, and isolates ESBL20150196 and ESBL20150300 (both obtained from Danish patients in 2015), each of which harbored a single 30-35-kb plasmid that, based on *in silico* mapping, was the result of a single major deletion that left only the RepFIB replicon (**Suppl. Fig. 3)**.

Next, we performed a SNP-based analysis of pAA-ST131 across all 57 *H*27 isolates with more than 70% coverage of the plasmid, with the four *H*22 clade EAEC isolates from the WHO collection used as an outgroup. This identified 197 SNPs within the ~25% of pAA-ST131 that was conserved across all isolates. Phylogenetic analysis based on these SNPs showed that pAA-ST131 is highly conserved among the *H*27 isolates (including the three with a truncated version of the plasmid), but differs considerably between the *H*27 and non-*H*27 isolates (**Suppl. Fig. 4**), strongly suggesting a single ancestral acquisition of the plasmid within the *H*27 clade, with subsequent partial deletions (to give the observed smaller variants).

Finally, we performed a BLAST analysis to screen the *H*27 isolates for classical ExPEC virulence genes (**Suppl. Table 3**). All *H*27 isolates carried the *sfa*, *iutA*, and *kpsM* II genes, which qualified them molecularly as ExPEC (*24*). In addition, they all contained the *chuA*, *fyuA*, and *yfcV* genes, which moreover qualified them molecularly as uropathogenic *E. coli* (UPEC) (*25*).

## Discussion

*E. coli* ST131 is a notorious MDR ExPEC lineage associated with both UTI and BSI (*1*). Little is known about which genes (whether promoting virulence or other phenotypes) make this clonal lineage so successful. ST131 strains have been shown to exhibit high levels of genomic plasticity including frequent recombination and plasmid flux, particularly involving IncF-type plasmids, facilitating spread of antibiotic resistance and virulence genes (*4, 26*). Indeed, ST131 lineages exhibit extensive variation with regard to acquired virulence genes (*2, 27*). Many typical ExPEC-associated virulence factors - including P fimbriae, hemolysins, and factors conferring increased serum survival and iron uptake - have been identified in ST131 isolates (*9, 10*). In contrast, virulence traits associated with diarrheagenic *E. coli* (DEC) pathotypes have thus far largely been absent from ST131 isolates *(10)*.

In developed countries, EAEC is known mostly as a cause of self-limiting diarrhea of mild to moderate severity. Indeed, long-term carriage of EAEC has been suggested to lead to colonization rather than infection (*11*). By contrast, in developing countries EAEC is a leading cause of childhood diarrhea (*11, 28*). The pathogenic potential of EAEC is underscored by its ability to cause major foodborne outbreaks of diarrheal disease (*19, 29, 30*). Intriguingly, recent studies have associated EAEC with UTI, suggesting that what classically has been regarded as a DEC pathotype may also qualify as ExPEC and cause both diarrhea and UTI (*31–33*).

Like other DEC pathotypes, EAEC has been shown to encompass diverse genetic lineages, reflecting a high level of phylogenetic heterogeneity (*16, 34*). Although until recently no extraintestinal EAEC ST131 had been documented, recent reports have described the occurrence in certain ST131 subclonal lineages of EAEC-specific traits in geographically distinct areas (*10, 13, 16*). ESBL-producing ST38 and other ST types of various serotypes have been indicated as emerging hybrid strains of UPEC and EAEC involved primarily in UTIs from Germany, the Netherlands and the United Kingdom (*35*). In a study of fecal and urine isolates from Danish patients in 1998-2000, we made the novel observation of EAEC ST131 of serotype O25 (*10*). Subsequently, another study documented the emergence of a *bla*_CTX-M-14_-containing EAEC ST131 O25:H4 strain in stool samples of diarrheic patients in Japan from 2003 onwards (*13*).

Our previous PFGE analysis of Danish ST131 O25 *E. coli* isolates from the WHO collection demonstrated that seven of the 12 identified EAEC strains were highly similar and appeared to have been part of an unrecognized UTI and diarrhea outbreak in Denmark in 1998 to 2000 (*10*). Although the genomic approach we used here to analyze the same isolates confirmed the suspected outbreak, it also yielded certain substantially different conclusions from the PFGE analysis, including identification of a second distinct cluster of EAEC urine isolates. These discrepancies between WGS SNP-based analysis and PFGE analysis among ST131 isolates correspond with previous findings (6).

Here we also document the current presence of a novel ESBL-EAEC ST131 sublineage as a cause of bacteremia in patients admitted to Danish hospitals. The isolates, as collected across Denmark over a 16-month period in 2014-2015, all carried *bla*_CTX-M-101_. Notably, to date the only other reports of CTX-M-101-positive *E. coli* are from China (*36–38*). Roer *et al*. used a genomic approach to establish that the Danish ESBL EAEC isolates were highly clonal, strongly suggesting a recent common source (*22*). Here, by analyzing sequential urine isolates from eight of the patients with the Danish outbreak-associated ESBL-EAEC ST131 bacteremia, we found that five patients had recurrent UTI caused by the outbreak strain. The time of paired urine sampling ranged from one week before to eight months following the initial blood sampling. The Danish ESBL-EAEC ST131 outbreak strain thus seems capable of persistently colonizing patients, resulting in occasional clinical manifestations. Less likely, the patients may have been continuously exposed from an external source.

Using the EnteroBase collection of ST131 genome sequences, we identified the global presence of *fimH*27-carrying EAEC ST131 isolates. These isolates originated from Africa, Asia, Europe and North America, and together with the Danish EAEC ST131 outbreak isolates spanned more than two decades. Strikingly, all the EAEC ST131 strains studied here proved to be clonal, i.e., to share a common ancestor. Moreover, they share a common virulence plasmid, designated pAA-ST131, which appears to have been acquired once in the *H*27 clade and transmitted vertically, although it has undergone recombination and deletion events in some sublineages. MinION-based sequencing of this plasmid in a representative ESBL-EAEC strain identified an array of classical plasmid-encoded EAEC-defining virulence genes, including those encoding AAFs and the global regulator of virulence AggR.

The acquisition of pAA plasmids by *E. coli* ST131 is interesting, considering that the ST131 lineage is thought of as a host generalist (*39*), whereas EAEC appears to be highly adapted to humans, without a natural animal reservoir (*40, 41*). However, the EAEC ST131 lineage described here is reminiscent of the UTI outbreak in Copenhagen in 1991 caused by an *E. coli* O78:H10 clonal group in that both fulfilled the molecular criteria for EAEC and contained multiple ExPEC virulence genes (*32*). That the O78:H10 outbreak strains’ EAEC-associated virulence factors were found to increase uropathogenicity (*33*) suggests that this may be true also for EAEC ST131 strains. The O78:H10 EAEC outbreak strain, which never was found outside the Capital Region of Denmark and was not known to have caused BSI. In contrast, ST131 is a highly successful pandemic clonal group, which makes the potential of future outbreaks of UTI and bacteremia caused by this novel strain, or other EAEC ST131 lineages, a cause for serious concern.

Despite the genetic evidence that all 25 *bla*_CTX-M-101_-containing Danish EAEC isolates were derived from a common source, based on the limited available epidemiological data we have been unable to establish a patient link or explain the spread of the lineage across regions. UTI is often caused by enteric *E. coli* that enter the urinary tract via the fecal-perineal-urethral route, and in some instances may have as their proximate external source food products or animals (*42*). Interestingly, the gender ratio among the 25 present cases (68% females, 32% males), differs significantly (*p* = 0.01) from that across the entire collection of 552 ESBL-producing ST131 bloodstream isolates from Danish hospitals in 2014-2015 (42% females, 58% males) (*22*). A similar overrepresentation of women was observed among cases during the 2011 multinational European outbreak caused by a novel multi-pathotype, Shiga-toxin-producing EAEC O104:H4 strain (*43*). Bean sprouts were found the most likely vehicle of infection, and the high proportion of female cases in that study was thought to be driven by the tendency of women to be more health conscious and perhaps suggestive of a food-related source for the Danish ESBL EAEC ST131 outbreak. Further screening, e.g. of fecal and urine samples from past and current patients who present with diarrhea or UTI, are warranted to determine the true clinical impact of the ESBL EAEC ST131 strain and to provide the demographic and epidemiological data needed to identify potential sources.

In conclusion, we hypothesize that acquisition of the pAA plasmid made this *H*27 ESBL EAEC ST131 lineage highly successful at persistently colonizing patients, thereby allowing it to occasionally cause UTIs and diarrhea. The presence of multiple ExPEC virulence factors - including P fimbriae, α-hemolysin, and Sat - in turn facilitates dissemination from the urinary tract to the bloodstream. Furthermore, we have demonstrated the ability of a non-*H*30 ST131 lineage to acquire EAEC-specific pAA virulence plasmids and to disseminate across multiple continents over the past two decades. An *H*27 EAEC ST131 strain that has also acquired the *bla*_CTX-M-101_ gene was causing bacteremia outbreaks in several geographic regions in Denmark, seemingly associated with recurrent infections. These findings emphasize the potential for different pathogens to evolve, thus potentially generating important new pathotypes that require continuous vigilance. They also illustrate the power of whole-genome sequencing to elucidate the historical and current molecular epidemiology and evolution of emerging pathogens of high public health importance.

## Materials and methods

### Genome data and sequencing

WGS was performed on a previously described international collection of 128 ST131 *E. coli* human isolates of serotype O25:H4 or O25:H- (1968 to 2011) from the WHO Collaborating Centre for Reference and Research on *Escherichia* and *Klebsiella* (www.ssi.dk) (*10*). DNA samples were prepared for multiplexed, paired-end sequencing using a combination of Illumina MiSeq and HiSeq (*6, 10*). We also included a published *E. coli* ST131 sequence dataset comprising 93 US and German isolates of human and animal origin (1967 to 2011) (*6*), and sequences from 27 ESBL-producing ST131 bloodstream isolates (carrying *bla*_CTX-M-101_) collected from Danish patients within the national surveillance program for antimicrobial resistance (DANMAP) for ESBL-producing *E. coli* (2014 - 2015) (*22*). Furthermore, all >3,500 *E. coli* ST131 genome data available at EnteroBase (http://enterobase.warwick.ac.uk, accessed November 15^th^ 2017) were analyzed for EAEC characteristics (presence for the *aggR* gene) and positive isolates were included. Finally, we sequenced and included 13 *E. coli* isolates obtained from urine from the source patients for eight of the ESBL-producing blood isolates. The sequences were analyzed using the Bacterial Analysis Platform (BAP) from Center for Genomic Epidemiology (*44*).

### Plasmid sequencing and analysis

Plasmid DNA was sequenced on both a MiSeq instrument (Illumina) and a MinION flow cell (Oxford Nanopore Technologies). The MiSeq library was made using the Nextera XT kit (Illumina) and sequencing was performed as a paired-end 250 bp run, yielding 372,720 reads with an average length of 237 bp. The MinION library was prepared using the Genomic Sequencing SQK-MAP006 kit and was sequenced on a FLO-MAP003 Early Access flow cell according to the manufacturer’s instruction. Fast5 read files were subjected to base calling via a two-direction workflow using Metrichor software (ONT), yielding 4,856 passed 2D read files. Mixed assembly was performed by combining MiSeq and MinION reads using the SPAdes assembler (v3.9.0). Finally, CLCbio Genomics Workbench (v9.5.2) was used for end trimming of the assembled plasmid and for final error correction by mapping trimmed MiSeq reads against the plasmid contig obtained after the mixed SPAdes assembly.

pAA-ST131 was annotated using RAST (*45*), with putative hypothetical genes curated manually using NCBI BLASTn and BLASTp searches. A BLASTn atlas of pAA-ST131 and other pAA virulence plasmids was constructed using BLAST Ring Image Generator v0.95 (BRIG) (*46*).

### Virulence genotyping

Isolates qualified molecularly as EAEC if positive for ≥1 of EAEC-associated putative virulence genes *aggR*, *aatA*, and *aaiC (10)*. Isolates were regarded as ExPEC if positive for ≥2 of *papA* and/or *papC* (P fimbriae), *sfa* and *foc* (S and F1C fimbriae), *afa* and *dra* (Dr-binding adhesins), *kspM* II (group 2 capsule), and *iutA* (aerobactin siderophore system) (*24*). Isolates were considered UPEC if positive for ≥2 of *chuA* (heme uptake), *fyuA* (yersiniabactin siderophore system), *vat* (vacuolating toxin), and *yfcV* (adhesin) (*25*).

### Identification of SNPs

SNPs in the core chromosome or plasmid, depending on analysis, were identified using the Northern Arizona SNP Pipeline (NASP) (*47*). Briefly, duplicate regions of the reference chromosome JJ1886 or pAA-ST131 plasmid (GenBank accession no. CP006784 and KY706108, respectively), were identified by aligning the reference against itself with NUCmer (*48*), followed by mapping of Illumina raw reads against the reference using the Burrows-Wheeler Aligner (BWA) (*49*) with identification of SNPs using GATK (*50*). Purging of recombinant region in the chromosome-based SNPs was performed using Gubbins v.2.2 (*51*).

### Phylogenetic analysis

Relatedness between isolates according to core genome SNPs was inferred using RAxML v8.2.10 using the GTRCAT model (*52*). Relatedness of the plasmids was inferred using using PHyML with Smart Model Selection (*53*) with tree searching using SPR and 100 bootstrap replicates.

### Plasmid profiling

Plasmids were purified as described by Kado and Liu (*54*) and visualized by separation on a 0.8% agarose gel electrophoresis and stained with GelRed (Biotium, Hayward, Ca, USA). *E. coli* strain 39R861, which contains four plasmids of sizes 147, 63, 36, and 7 kb (*55, 56*), served as a size marker.

### Statistical Analysis

Comparisons of categorical variables were tested using Pearson’s χ^2^ test with Yates’ continuity correction, using R v3.3.2 statistical software (https://www.r-project.org). The criterion for statistical significance was p<0.05.

### Accession of Sequence data

The accession numbers for the Illumina sequences generated from the 134 *E. coli* ST131 isolates presented in this study are available in the European Nucleotide Archive (ENA; https://www.ebi.ac.uk/ena) under the following accession numbers: PRJEB27194. Sequences can also be located in the ENA using the following study summary: “Emergence of enteroaggregative *Escherichia coli* within the ST131 lineage as a cause of extraintestinal infections”. The sequence of the pAA-ST131 plasmid has been deposited in GenBank under accession number KY706108.

## Acknowledgements

We thank Michala T. Sørensen at Statens Serum Institut for performing the plasmid gel electrophoresis. We also thank Karin Sixhøj Pedersen at Statens Serum Institut for her excellent WGS work.

---

**Suppl. Fig. 1.** Rooted phylogenetic tree of ST131 *E. coli* strains from the WHO collection (n=121) and the US/German collection (n=93). Isolates fulfilling the molecular criteria for EAEC are marked with an asterisk.

**Suppl. Fig. 2.** Midpoint-rooted phylogeny based on 58 SNPs of paired blood (red) and urine (yellow) isolates from eight Danish cases as well as all remaining Danish ST131 EAEC *H*27 *bla*_CTX-M-101_-positive isolates. The analysis depicts the distinct relatedness of five corresponding blood and urine isolates (Cases 1, 4, 5, 7, and 8), whereas the data is inconclusive for the remaining isolates (Cases 2, 3 and 6).

**Suppl. Fig. 3.** Plasmid profiles of representative *fimH*27-carrying ST131 isolates either containing *bla*_CTX-M-101_ or from the WHO collection. Plasmid profiles with *E. coli* 39R861 as a marker (147, 63, 36 and 7kb) are included in lane 1. All ten *fimH*27 strains harbored only a single a plasmid. In seven of the strains, this plasmid was 140-145 kb in size, whereas a slightly smaller size plasmid was present in WHO isolate C1883-99. ESBL20150196 and ESBL20150300 both harbored a significantly smaller plasmid of approximately 30 kb.

**Suppl. Fig. 4.** Phylogenetic tree of pAA from ST131 *H*27 isolates with more than 70% coverage of pAA-ST131 from representative Danish isolate ESBL20150001. Three of the *H*22/*H*298 EAEC isolates from the WHO collection (C156-00, C168-00 and C180-00) were included as an outgroup. A total of 197 SNPs was identified equivalent to approximately 25% of pAA-ST131.

## References

1. Rogers BA, Sidjabat HE, Paterson DL. 2011. Escherichia coli O25b-ST131: a pandemic, multiresistant, community-associated strain. J Antimicrob Chemother 66: 1–14.

2. Johnson JR, Johnston B, Clabots C, Kuskowski MA, Castanheira M. 2010. Escherichia coli sequence type ST131 as the major cause of serious multidrug-resistant E. coli infections in the United States. Clin Infect Dis 51: 286–294.

3. Banerjee R, Johnson JR. 2014. A new clone sweeps clean: the enigmatic emergence of Escherichia coli sequence type 131. Antimicrob Agents Chemother 58: 4997–5004.

4. Stoesser N, Sheppard AE, Pankhurst L, De Maio N, Moore CE, Sebra R, Turner P, Anson LW, Kasarskis A, Batty EM, Kos V, Wilson DJ, Phetsouvanh R, Wyllie D, Sokurenko E, Manges AR, Johnson TJ, Price LB, Petro TE, Johnson JR, Didelot X, Walker AS, Crook DW, Modernizing Medical Microbiology Informatics Group (MMMIG). 2016. Evolutionary History of the Global Emergence of the Escherichia coli Epidemic Clone ST131. MBio 7: e02162–15.

5. Ben Zakour NL, Alsheikh-Hussain AS, Ashcroft MM, Khanh Nhu NT, Roberts LW, Stanton-Cook M, Schembri MA, Beatson SA. 2016. Sequential Acquisition of Virulence and Fluoroquinolone Resistance Has Shaped the Evolution of Escherichia coli ST131. MBio 7: e00347–00316.

6. Price LB, Johnson JR, Aziz M, Clabots C, Johnston B, Tchesnokova V, Nordstrom L, Billig M, Chattopadhyay S, Stegger M, Andersen PS, Pearson T, Riddell K, Rogers P, Scholes D, Kahl B, Keim P, Sokurenko EV. 2013. The epidemic of extended-spectrum-β-lactamase-producing Escherichia coli ST131 is driven by a single highly pathogenic subclone, H30-Rx. MBio 4: e00377–00313.

7. Matsumura Y, Johnson JR, Yamamoto M, Nagao M, Tanaka M, Takakura S, Ichiyama S, Kyoto-Shiga Clinical Microbiology Study Group. 2015. CTX-M-27- and CTX-M-14-producing, ciprofloxacin-resistant Escherichia coli of the H30 subclonal group within ST131 drive a Japanese regional ESBL epidemic. J Antimicrob Chemother 70: 1639–1649.

8. Abe CM, Salvador FA, Falsetti IN, Vieira MA, Blanco J, Blanco JE, Blanco M, Machado AM, Elias WP, Hernandes RT, Gomes TA. 2008. Uropathogenic Escherichia coli (UPEC) strains may carry virulence properties of diarrhoeagenic E. coli. FEMS Immunol Med Microbiol 52: 397–406.

9. Blanco J, Mora A, Mamani R, Lopez C, Blanco M, Dahbi G, Herrera A, Marzoa J, Fernandez V, de la Cruz F, Martinez-Martinez L, Alonso MP, Nicolas-Chanoine MH, Johnson JR, Johnston B, Lopez-Cerero L, Pascual A, Rodriguez-Bano J, Spanish Group for Nosocomial Infections (GEIH). 2013. Four main virotypes among extended-spectrum-β-lactamase-producing isolates of Escherichia coli O25b:H4-B2-ST131: bacterial, epidemiological, and clinical characteristics. J Clin Microbiol 51: 3358–3367.

10. Olesen B, Frimodt-Møller J, Leihof RF, Struve C, Johnston B, Hansen DS, Scheutz F, Krogfelt KA, Kuskowski MA, Clabots C, Johnson JR. 2014. Temporal trends in antimicrobial resistance and virulence-associated traits within the Escherichia coli sequence type 131 clonal group and its H30 and H30-Rx subclones, 1968 to 2012. Antimicrob Agents Chemother 58: 6886–6895.

11. Hebbelstrup Jensen B, Olsen KE, Struve C, Krogfelt KA, Petersen AM. 2014. Epidemiology and clinical manifestations of enteroaggregative Escherichia coli. Clin Microbiol Rev 27: 614–630.

12. Rasko DA, Webster DR, Sahl JW, Bashir A, Boisen N, Scheutz F, Paxinos EE, Sebra R, Chin CS, Iliopoulos D, Klammer A, Peluso P, Lee L, Kislyuk AO, Bullard J, Kasarskis A, Wang S, Eid J, Rank D, Redman JC, Steyert SR, Frimodt-Møller J, Struve C, Petersen AM, Krogfelt KA, Nataro JP, Schadt EE, Waldo MK. 2011. Origins of the E. coli strain causing an outbreak of hemolytic-uremic syndrome in Germany. N Engl J Med 365: 709–717.

13. Imuta N, Ooka T, Seto K, Kawahara R, Koriyama T, Kojyo T, Iguchi A, Tokuda K, Kawamura H, Yoshiie K, Ogura Y, Hayashi T, Nishi J. 2016. Phylogenetic Analysis of Enteroaggregative Escherichia coli (EAEC) Isolates from Japan Reveals Emergence of CTX-M-14-Producing EAEC O25:H4 Clones Related to Sequence Type 131. J Clin Microbiol 54: 2128–2134.

14. Guiral E, Mendez-Arancibia E, Soto SM, Salvador P, Fabrega A, Gascon J, Vila J. 2011. CTX-M-15- producing enteroaggregative Escherichia coli as cause of travelers’ diarrhea. Emerg Infect Dis 17: 1950–1953.

15. Amaya E, Reyes D, Vilchez S, Paniagua M, Möllby R, Nord CE, Weintraub A. 2011. Antibiotic resistance patterns of intestinal Escherichia coli isolates from Nicaraguan children. J Med Microbiol 60: 216–222.

16. Zhang R, Gu DX, Huang YL, Chan EW, Chen GX, Chen S. 2016. Comparative genetic characterization of Enteroaggregative Escherichia coli strains recovered from clinical and non-clinical settings. Sci Rep 6: 24321.

17. Nataro JP. 2005. Enteroaggregative Escherichia coli pathogenesis. Curr Opin Gastroenterol 21: 4–8.

18. Santiago AE, Ruiz-Perez F, Jo NY, Vijayakumar V, Gong MQ, Nataro JP. 2014. A large family of antivirulence regulators modulates the effects of transcriptional activators in Gram-negative pathogenic bacteria. PLoS Pathog 10: e1004153.

19. Boisen N, Scheutz F, Rasko DA, Redman JC, Persson S, Simon J, Kotloff KL, Levine MM, Sow S, Tamboura B, Toure A, Malle A, Malle D, Panchalingam S, Krogfelt KA, Nataro JP. 2012. Genomic characterization of enteroaggregative Escherichia coli from children in Mali. J Infect Dis 205: 431–444.

20. Jønsson R, Struve C, Boll EJ, Boisen N, Joensen KG, Sørensen CA, Jensen BH, Scheutz F, Jenssen H, Krogfelt KA. 2015. Novel aggregative adherence fimbria variant of enteroaggregative Escherichia coli. Infect Immun 83: 1396–1405.

21. Morin N, Santiago AE, Ernst RK, Guillot SJ, Nataro JP. 2013. Characterization of the AggR Regulon in Enteroaggregative Escherichia coli. Infect Immun 81: 122–132.

22. Roer L, Hansen F, Thomsen MCF, Knudsen JD, Hansen DS, Wang M, Samulioniene J, Justesen US, Røder BL, Schumacher H, Østergaard C, Andersen LP, Dzajic E, Søndergaard TS, Stegger M, Hammerum AM, Hasman H. 2017. WGS-based surveillance of third-generation cephalosporin-resistant Escherichia coli from bloodstream infections in Denmark. J Antimicrob Chemother 72: 1922–1929.

23. Johnson TJ, Nolan LK. 2009. Pathogenomics of the virulence plasmids of Escherichia coli. Microbiol Mol Biol Rev 73: 750–774.

24. Johnson JR, Murray AC, Gajewski A, Sullivan M, Snippes P, Kuskowski MA, Smith KE. 2003. Isolation and molecular characterization of nalidixic acid-resistant extraintestinal pathogenic Escherichia coli from retail chicken products. Antimicrob Agents Chemother 47: 2161–2168.

25. Spurbeck RR, Dinh PC Jr, Walk ST, Stapleton AE, Hooton TM, Nolan LK, Kim KS, Johnson JR, Mobley HL. 2012. Escherichia coli isolates that carry vat, fyuA, chuA, and yfcV efficiently colonize the urinary tract. Infect Immun 80: 4115–4122.

26. Lanza VF, de Toro M, Garcillan-Barcia MP, Mora A, Blanco J, Coque TM, de la Cruz F. 2014. Plasmid flux in Escherichia coli ST131 sublineages, analyzed by plasmid constellation network (PLACNET), a new method for plasmid reconstruction from whole genome sequences. PLoS Genet 10: e1004766.

27. Coelho A, Mora A, Mamani R, Lopez C, Gonzalez-Lopez JJ, Larrosa MN, Quintero-Zarate JN, Dahbi G, Herrera A, Blanco JE, Blanco M, Alonso MP, Prats G, Blanco J. 2011. Spread of Escherichia coli O25b:H4-B2-ST131 producing CTX-M-15 and SHV-12 with high virulence gene content in Barcelona (Spain). J Antimicrob Chemother 66: 517–526.

28. Okhuysen PC, Dupont HL. 2010. Enteroaggregative Escherichia coli (EAEC): a cause of acute and persistent diarrhea of worldwide importance. J Infect Dis 202: 503–505.

29. Itoh Y, Nagano I, Kunishima M, Ezaki T. 1997. Laboratory investigation of enteroaggregative Escherichia coli O untypeable:H10 associated with a massive outbreak of gastrointestinal illness. J Clin Microbiol 35: 2546–2550.

30. Dallman TJ, Chattaway MA, Cowley LA, Doumith M, Tewolde R, Wooldridge DJ, Underwood A, Ready D, Wain J, Foster K, Grant KA, Jenkins C. 2014. An investigation of the diversity of strains of enteroaggregative Escherichia coli isolated from cases associated with a large multi-pathogen foodborne outbreak in the UK. PLoS One 9: e98103.

31. Park HK, Jung YJ, Chae HC, Shin YJ, Woo SY, Park HS, Lee SJ. 2009. Comparison of Escherichia coli uropathogenic genes (kps, usp and ireA) and enteroaggregative genes (aggR and aap) via multiplex polymerase chain reaction from suprapubic urine specimens of young children with fever. Scand J Urol Nephrol 43: 51–57.

32. Olesen B, Scheutz F, Andersen RL, Menard M, Boisen N, Johnston B, Hansen DS, Krogfelt KA, Nataro JP, Johnson JR. 2012. Enteroaggregative Escherichia coli O78:H10 -- the Cause of an Outbreak of Urinary Tract Infection. J Clin Microbiol 50: 3703–11.

33. Boll EJ, Struve C, Boisen N, Olesen B, Stahlhut SG, Krogfelt KA. 2013. Role of enteroaggregative Escherichia coli virulence factors in uropathogenesis. Infect Immun 81: 1164–1171.

34. Okeke IN, Wallace-Gadsden F, Simons HR, Matthews N, Labar AS, Hwang J, Wain J. 2010. Multi-locus sequence typing of enteroaggregative Escherichia coli isolates from Nigerian children uncovers multiple lineages. PLoS One 5: e14093.

35. Chattaway MA, Jenkins C, Ciesielczuk H, Day M, DoNascimento V, Day M, Rodiguez I, van Essen-Zandbergen A, Schink AK, Wu G, Threlfall J, Woodward MJ, Coldham N, Kadlec K, Schwarz S, Dierikx C, Guerra B, Helmuth R, Mevius D, Woodford N, Wain J. 2014. Evidence of evolving extraintestinal enteroaggregative Escherichia coli ST38 clone. Emerg Infect Dis 20: 1935–1937.

36. Zhang J, Zheng B, Zhao L, Wei Z, Ji J, Li L, Xiao Y. 2014. Nationwide high prevalence of CTX-M and an increase of CTX-M-55 in Escherichia coli isolated from patients with community-onset infections in Chinese county hospitals. BMC Infect Dis 14: 659.

37. Xia S, Fan X, Huang Z, Xia L, Xiao M, Chen R, Xu Y, Zhuo C. 2014. Dominance of CTX-M-type extended-spectrum β-lactamase (ESBL)-producing Escherichia coli isolated from patients with community-onset and hospital-onset infection in China. PLoS One 9: e100707.

38. Tong P, Sun Y, Ji X, Du X, Guo X, Liu J, Zhu L, Zhou B, Zhou W, Liu G, Feng S. 2015. Characterization of antimicrobial resistance and extended-spectrum β-lactamase genes in Escherichia coli isolated from chickens. Foodborne Pathog Dis 12: 345–352.

39. McNally A, Oren Y, Kelly D, Pascoe B, Dunn S, Sreecharan T, Vehkala M, Välimäki N, Prentice MB, Ashour A, Avram O, Pupko T, Dobrindt U, Literak I, Guenther S, Schaufler K, Wieler LH, Zhiyong Z, Sheppard SK, McInerney JO, Corander J. 2016. Combined Analysis of Variation in Core, Accessory and Regulatory Genome Regions Provides a Super-Resolution View into the Evolution of Bacterial Populations. PLoS Genet 12: e1006280.

40. Trung NV, Nhung HN, Carrique-Mas JJ, Mai HH, Tuyen HT, Campbell J, Nhung NT, Van Minh P, Wagenaar JA, Mai NT, Hieu TQ, Schultsz C, Hoa NT. 2016. Colonization of Enteroaggregative Escherichia coli and Shiga toxin-producing Escherichia coli in chickens and humans in southern Vietnam. BMC Microbiol 16: 208.

41. Puño-Sarmiento J, Medeiros L, Chiconi C, Martins F, Pelayo J, Rocha S, Blanco J, Blanco M, Zanutto M, Kobayashi R, Nakazato G. 2013. Detection of diarrheagenic Escherichia coli strains isolated from dogs and cats in Brazil. Vet Microbiol 166: 676–680.

42. Mitsumori K, Terai A, Yamamoto S, Yoshida O. 1997. Virulence characteristics and DNA fingerprints of Escherichia coli isolated from women with acute uncomplicated pyelonephritis. J Urol 158: 2329–2332.

43. Frank F, Werber D, Cramer JP, Askar M, Faber M, an der Heiden M, Bernard H, Fruth A, Prager R, Spode A, Wadl M, Zoufaly A, Jordan S, Kemper MJ, Follin P, Müller L, King LA, Rosner B, Buchholz U, Stark K, Krause G, HUS Investigation Team. 2011. Epidemic profile of Shiga-toxin-producing Escherichia coli O104:H4 outbreak in Germany. N Engl J Med 365: 1771–1780.

44. Thomsen MC, Ahrenfeldt J, Cisneros JL, Jurtz V, Larsen MV, Hasman H, Aarestrup FM, Lund O. 2016. A Bacterial Analysis Platform: An Integrated System for Analysing Bacterial Whole Genome Sequencing Data for Clinical Diagnostics and Surveillance. PLoS One 11: e0157718.

45. Aziz RK, Bartels D, Best AA, DeJongh M, Disz T, Edwards RA, Formsma K, Gerdes S, Glass EM, Kubal M, Meyer F, Olsen GJ, Olson R, Osterman AL, Overbeek RA, McNeil LK, Paarmann D, Parczian T, Parrello B, Pusch GD, Reich C, Stevens R, Vassieva O, Vonstein V, Wilke A, Zagnitko O. 2008. The RAST Server: rapid annotations using subsystems technology. BMC Genomics 9: 75.

46. Alikhan NF, Petty NK, Ben Zakour NL, Beatson SA. 2011. BLAST Ring Image Generator (BRIG): simple prokaryote genome comparisons. BMC Genomics 12: 402.

47. Sahl JW, Lemmer D, Travis J, Schupp JM, Gillece JD, Aziz M, Driebe EM, Drees KP, Hicks ND, Williamson CHD, Hepp CM, Smith DE, Roe C, Engelthaler DM, Wagner DM, Keim P. 2016. NASP: an accurate, rapid method for the identification of SNPs in WGS datasets that supports flexible input and output formats. Microb Genom 2: e000074.

48. Delcher AL, Phillippy A, Carlton J, Salzberg SL. 2002. Fast algorithms for large-scale genome alignment and comparison. Nucleic Acids Res 30: 2478–2483.

49. Li H, Durbin R. 2009 Fast and accurate short read alignment with Burrows-Wheeler transform. Bioinformatics 25: 1754–1760.

50. McKenna A, Hanna M, Banks E, Sivachenko A, Cibulskis K, Kernytsky A, Garimella K, Altshuler D, Gabriel S, Daly M, DePristo MA. 2010. The Genome Analysis Toolkit: a MapReduce framework for analyzing next-generation DNA sequencing data. Genome Res 20: 1297–1303.

51. Croucher NJ, Page AJ, Connor TR, Delaney AJ, Keane JA, Bentley SD, Parkhill J, Harris SR. 2015. Rapid phylogenetic analysis of large samples of recombinant bacterial whole genome sequences using Gubbins. Nucleic Acids Res 43: e15.

52. Stamatakis A. 2014. RAxML version 8: a tool for phylogenetic analysis and post-analysis of large phylogenies. Bioinformatics 30: 1312–1313.

53. Lefort V, Longueville JE, Gascuel O. 2017. SMS: Smart Model Selection in PhyML. Mol Biol Evol 34: 2422–2424.

54. Kado CI, Liu ST. 1981. Rapid procedure for detection and isolation of large and small plasmids. J Bacteriol 145: 1365–1373.

55. Threlfall EJ, Rowe B, Ferguson JL, Ward LR. 1986. Characterization of plasmids conferring resistance to gentamicin and apramycin in strains of Salmonella typhimurium phage type 204c isolated in Britain. J Hyg (Lond) 97: 419–426.

56. Schjørring S, Struve C, Krogfelt KA. 2008. Transfer of antimicrobial resistance plasmids from Klebsiella pneumoniae to Escherichia coli in the mouse intestine. J Antimicrob Chemother 62: 1086–1093.

